# Deciphering interplay between biology and physics: finite element method-implemented vertex organoid model raises the challenge

**DOI:** 10.1101/2023.05.15.540870

**Authors:** J. Laussu, D. Michel, L. Magne, S. Segonds, S. Marguet, D. Hamel, M. Quaranta-Nicaise, F. Barreau, E. Mas, V. Velay, F. Bugarin, A. Ferrand

## Abstract

Understanding the intertwining of biology and mechanics in tissue architecture is a challenging issue, especially when it comes to the 3D tissue organization. Addressing this challenge requires both a biological model allowing multiscale observations from the cell to the tissue, and theoretical and computational approaches allowing the generation of a synthetic model, relevant to the biological model, and allowing access to the mechanical constraints experienced by the tissue.

Here, using human colon epithelium monolayer organoid as biological model, and combining vertex and FEM approaches, we generated a comprehensive elastic finite element model of the human colon organoid and demonstrated its flexibility. This FEM model provides a basis for relating cell shape, tissue deformation, and strain at the cellular level due to imposed stresses.

In conclusion, we demonstrated that the combination of vertex and FEM approaches allows for better modeling of the alteration of organoid morphology over time and better assessment of the mechanical cues involved in establishing the architecture of the human colon epithelium.

## INTRODUCTION

Understanding the interplay between biology and mechanics involved in tissue architecture is challenging, especially when it comes to 3D tissue organization. It requires an appropriate biological model, allowing multi-scale observations from cellular resolution to global tissue architecture, and pertinent theoretical and computational approaches to generate a synthetic model being the most relevant to the biological model, and allowing accessing the mechanical stresses undergone by the tissue.

A useful biological model to address this interplay is the organoid, especially when recreating a simple monolayer epithelial tissue. Organoids are simplified versions of organs and tissues, established from either adult stem cells, embryonic stem cells (ESCs) or induced pluripotent stem cells (iPSCs), owing to their self-renewal and differentiation capacities. Cultured in three dimensions (3D) in supporting matrix, organoids are autonomous, self-organized, recapitulate the original tissue architecture and at least one tissue function^1–8^. Moreover, the *in vitro* culture facilitates their live observation by microscopy allowing to access relatively high temporal resolution during the follow-up of the architecture establishment. Therefore, their ability to integrate complex cell behaviors in a relatively short timeline make this model a perfect tool to study how biology and physics of the cell cooperate to establish tissue architecture.

Regarding the apprehension of the physics stress, theoretical and computational approaches inspired by physics and combined to experimental observations of this biological model have been developed^9^. Recent synthetic models focus on 3D processes occurring in morphogenetic events^10–12^. Among them, 3D vertex models are well suited for biological systems since taking into account macroscopic mechanical characteristics while allowing cell identity representation^13^. Indeed, vertex models can be seen as geometrical description of the biological system where each cell is represented by polygons described by vertices connected with edges, and a discretization of this geometry using basic elements upon which the mechanical equations are solved. This ability to represent each cell individually allows to study tissue organization. 3D vertex model can be considered as a primitive Finite Element Method (FEM) model to access mechanical properties^14^. FEM utilizes volumetric meshes to compute internal stresses and forces throughout the volume of an object. Overall, FEM is a well-established numerical analysis method employed in mechanics to model deformation of complex structures for which obtaining an analytical solution is difficult. Moreover, FEM is well suited to model the effect of deformation due to local constraints. It uses large collections of predefined elements to approximate the complete partition by polynomial functions on each element. Therefore, FEM, proposing a discrete solution of a continuous problem, has been employed for solid tissue modeling^15^.

Here, using the monolayer human colon epithelium organoid as biological model, and combining vertex and FEM approaches, we generated an innovative *in silico* human colon organoid model giving access to the interplay between biology and mechanics.

Using an approach combining real-time fluorescence imaging and 2D and 3D image analysis, we extracted biological parameters characterizing the organoid biological model during its architecture establishment. Then, their implementations into an ‘organoid’ vertex model allowed us to observe the limits of this vertex method by comparing the data obtained on the biological organoid and the *in silico* vertex model. Finally, to obtain a more relevant i*n silico* organoid model and to understand the impact of the physics involved in the dynamic morphological changes contributing to the establishment of the colorectal organoid architecture, we generated an innovative specific FEM model. Analysis of the obtained results demonstrates how the flexibility of our FEM model can provide a basis for linking cell shape, tissue deformation and stresses at the cellular level due to imposed constraints.

In conclusion, we demonstrated that combining vertex and FEM approaches allow a better modeling of the organoid morphology alteration over time and a better evaluation of the mechanical clues involved in the establishment of the human colon epithelium architecture.

## RESULTS

### Extraction of the biological parameters from the human colon organoids

Morphology being one of the principal features defining intestinal monolayer epithelial organoid^16^, we used the volumetric vertex model of organoids combined with FEM in order to simulate the human colon organoid architecture over time, using simple physical law and material properties.

The experimental efforts to understand this monolayer epithelium homeostasis have largely focused on the study of cell proliferation, cell differentiation and cell death regulation^17,18^. However other parameters associated to tissue architecture, such as cell shape regulation^19,20^, or mechanical stress, namely osmotic pressure^21^, external solicitation^22^ and active contraction^23^, have been proved involved in epithelial tissue shaping. Deciphering the individual contributions of these intertwined cell events into the intestinal tissue architecture establishment necessitates producing a powerful computational model gathering biology, physics, and geometry.

Our first objective was extracting architectural/geometrical parameters from the human colon organoids at different stages of tissue architecture establishment. As described^24^, human colon organoids are established from isolated crypts extracted from patient’s colon biopsies. Crypts are cultivated in Matrigel matrix to obtain colon organoids (Fig. 1A). Over the culture, we followed the establishment of the organoid architecture and acquired stack images in order to reconstruct the organoid 3D morphology and access its parameters at cellular resolution. The mean size of the imaged organoids were about 350 µm of diameter, for a mean number of cells equivalent at 390 cells per organoid (Table 1). As highlighted by staining for nuclei (DRAQ5 or SpyDNA), cell membrane proteins (EpCAM) or actin-myosin cytoskeleton (visualized thanks to actin staining with Phalloidin or FastActin), colon organoids display different morphologies along their culture. They evolve from a ‘cystic’ morphology (Fig. 1C, T0) with a nearly spherical flat epithelium (flattened nuclei, elongated flattened cells) and a central lumen (L), to a ‘columnar’ morphology (Fig. 1C, T0 + 180 minutes) displaying a thicker polarized epithelium (nuclei at basal position, decrease of apical and basal poles of the cells associated to lateral sides elongation, and swelling of the apical membrane), reduced central lumen (L) and deformations leading to a less spherical global organoid morphology (Fig. 1B, 1C, 1D). An in-between step (Fig. 1C, T0 + 90 minutes) is named here ‘intermediate’ morphology (Fig. 1C & 1D).

**Figure 1.**
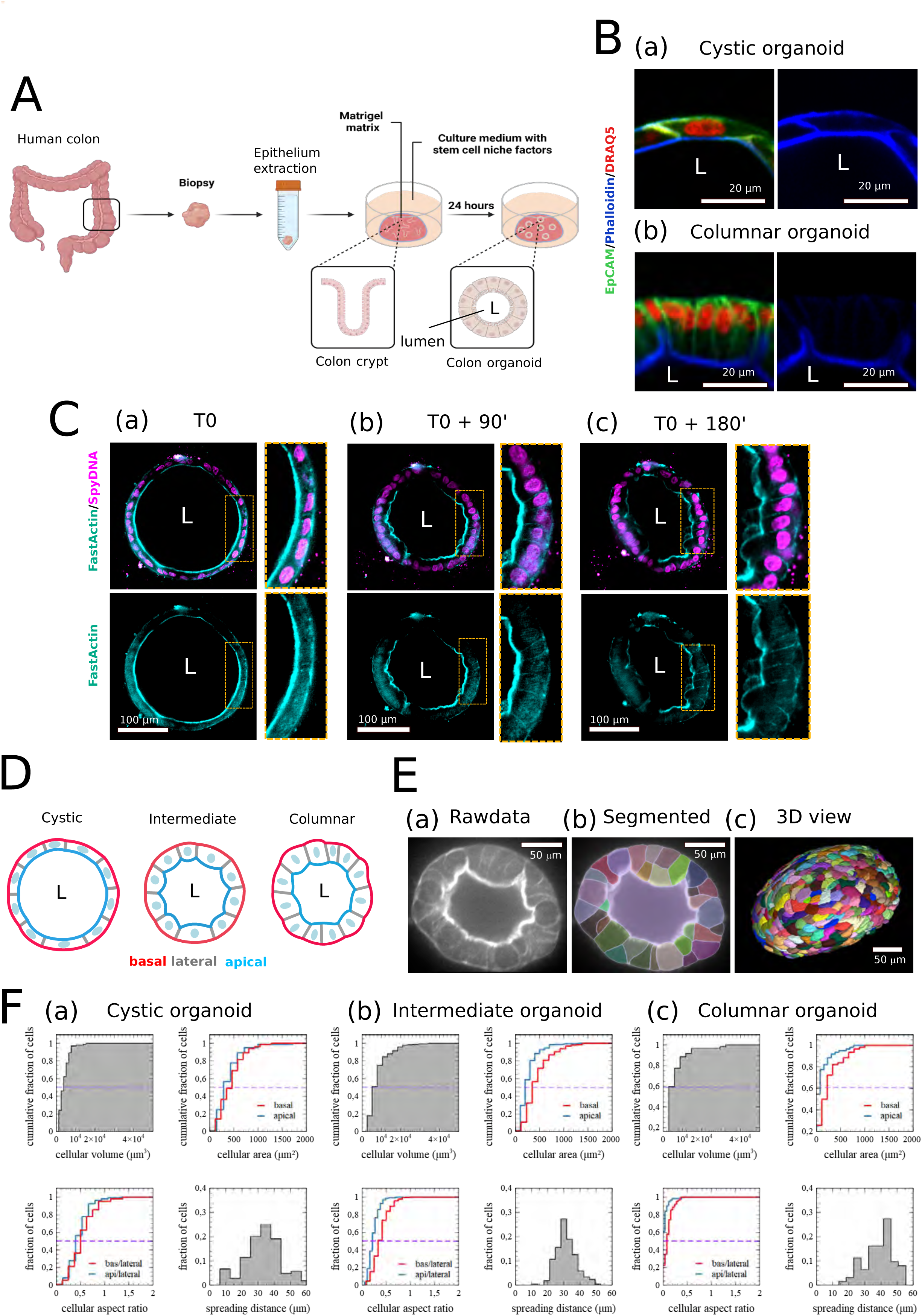
A/ Schema representing the experimental process to obtain colon organoid from human biopsies after isolation and culture in Matrigel matrix of the crypts. Illustration was created with BioRender.com. B/ High magnification at cellular scale of a confocal snapshot imaged on a fixed colon organoid with phalloidin (blue) immunostaining, EpCAM (green) immunostaining, and nuclear Draq5 (red) staining illustrating (a) a cystic organoid or (b) a columnar organoid after 15 days of culture in Matrigel. C/ Growth pattern of human intestinal organoids in Matrigel on a 3-hour timelapse analysis actin FastActin dye (cyan) and nuclei SpyDNA dye (purple). After deflation and a loss of organoid sphericity (T0 + 90’) (b) in a short time window, the apical membrane become more and more irregular, cells nuclei orientation changes to parallel to the apical border to perpendicular to it (T0 + 180’) (c). D/ Schema illustrating the global and cellular deformation characterizing the Cystic, Intermediate and Columnar morphology of the living organoids observed before. E/ Presentation of the protocol used to quantify organoid morphology. (a) Based on FastActin imaged on confocal microscopy, (b) cellular compartments are segmented using machine learning based approach (Cellpose) first in 2D and secondly stitches and interpolated in 3D; lumen is manually segmented using Napari in order to define lumen/apical interface. 3D exploration and manual correction of another bigger segmented organoid performed using Morphonet or Napari(c). F/ Morphology description card of three different organoids resumed with four different distribution plots for the cellular volume, basal (red) and apical (blue) surface area, basal/lateral (bas/lateral, red) and apical/lateral (api/lateral, blue) aspect ratio and spreading distance distribution for one cystic organoid (a), one intermediate organoid (b) and one columnar organoid (c).

**Table 1:** Parameters extracted from the live observation and analysis of the biological human colon organoid model.

| Morphology | Organoid ID | Diameter ($\mu\text{m}$) | Cell number (estimated) | Cell number (analyzed) | Thickness ratio |
| --- | --- | --- | --- | --- | --- |
| Cystic | 1 | 513 | 387 | 194 | 0.084 |
| Cystic | 2 | 572 | 412 | 226 | 0.055 |
| Cystic | 3 | 290 | 276 | 173 | 0.085 |
| Intermediate | 4 | 230 | 369 | 205 | 0.275 |
| Intermediate | 5 | 178 | 403 | 345 | 0.194 |
| Intermediate | 6 | 277 | 520 | 312 | 0.283 |
| Columnar | 7 | 265 | 387 | 209 | 0.512 |
| Columnar | 8 | 384 | 241 | 168 | 0.41 |
| Columnar | 9 | 474 | 518 | 363 | 0.342 |

Cells being the morphological units of the tissues, we can consider these global organoid morphology changes linked to architectural alterations at cellular level. Thus, we first aimed to extract individual cell information. Thanks to the actin-myosin cytoskeleton organization and based on its fluorescent staining (Fig. 1Ea), we used Cellpose 2.0^25,26^ to perform a 2D segmentation followed by a 3D organoid reconstruction (Fig. 1Ec). This approach, on cystic, intermediate and columnar morphological stages, permitted to extract for each cell its volume, apical and basal area, lateral area (cell-cell surface) allowing to evaluate the cellular aspect ratio (Fig. 1F). Moreover, to integrate topological information, we calculated a spreading distance corresponding to the mean distance between a cell centroid and the centroids of the five nearest neighboring cells. Five being chosen because it corresponds to the minimal number of neighboring cells for one single cell within the organoid 3D reconstruction^27^.

Based on these data, we then developed an *in silico* model of the intestinal organoid.

### Generation of vertex model for human colon organoid

In our organoid vertex model, each cell is a polygon mesh, a collection of vertices, faces and edges (Fig. 2A). Vertices are the ensemble of edges nodes. Faces are the external structures of the cells, for both the cell-cell and the cell-exterior interfaces. Edges are segments at the perimeter of the polygonal faces. Thus, to individualize each cell within the global structure, vertices, faces and edges define and are dependent on a cell identity.

**Figure 2.**
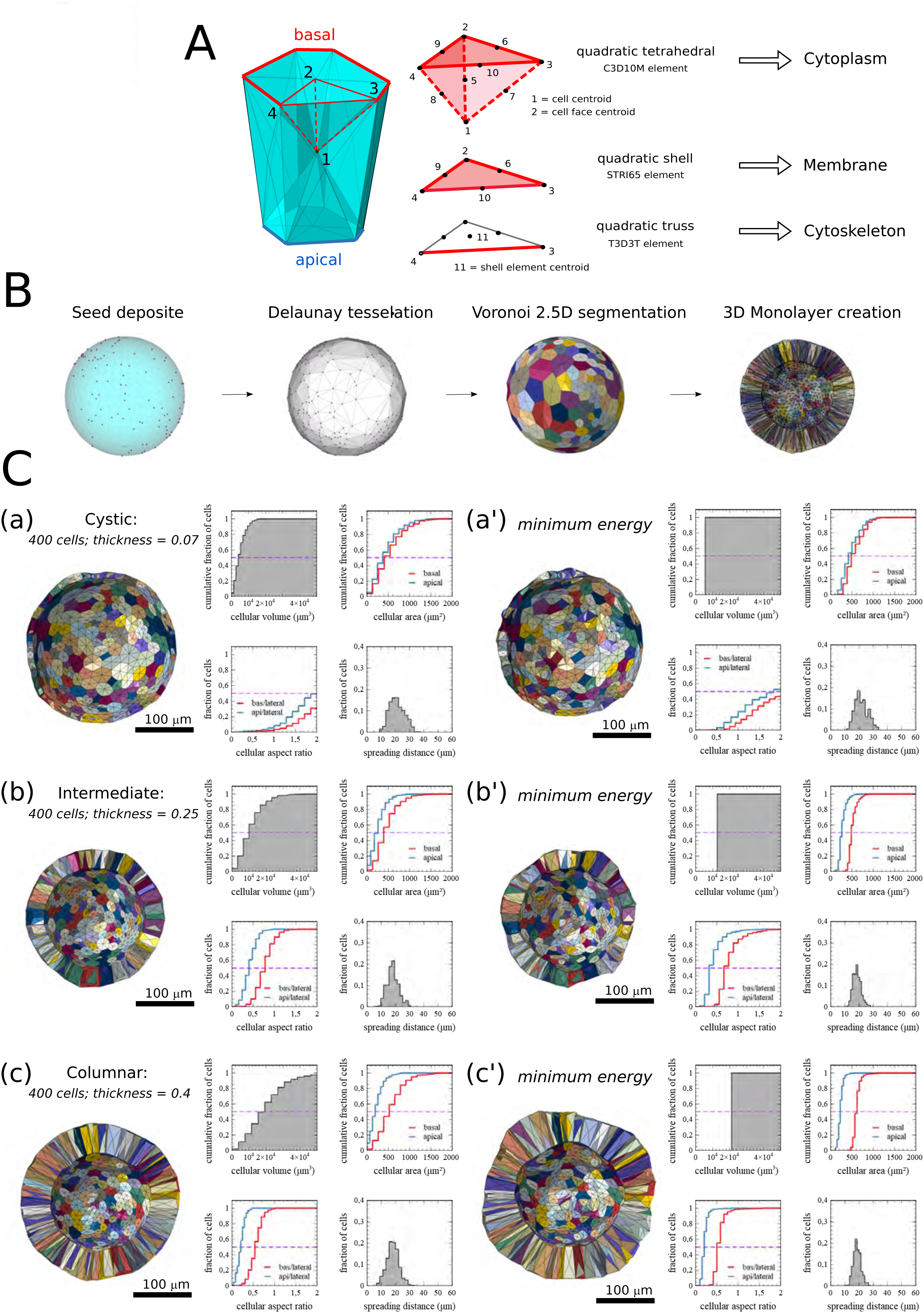
A/ Description of the subcellular element model. Cell is composed of three different types of elements: Quadratic tetrahedral elements for the cytoplasm volume discretization using C3D10M predefined Abaqus© element, quadratic Shell elements for the membrane discretization using STRI65 predefined Abaqus© element and quadratic truss for the cortex discretization using T3D3T predefined Abaqus© element contouring for the apical, basal and lateral surface of the cells to draw a ‘ring like’ structure. B/ *In silico* organoid are generated after four successive steps: seeds are non-uniformly distributed using random function at the surface of a normalized sphere. A Delaunay 3D create convex hull of the surface and a Voronoi 3D step segment cellular distinct area. Finally, the) creation of a thickness occurs using the dilation of the sphere using a predefined “thickness ratio” relative to the radius of the normal sphere. C/ Using previously published 3D Active Vertex Model (AVM) resolution to model the three ‘cystic’ (a, a’), ‘intermediate’ (b, b’) and ‘columnar’ (c, c’) organoids shape evolution. This AVM is a rheological model using gradient descent energy minimization to calculate the equilibrium state of the *in silico* organoid (a’, b’, c’) ^33^.

To construct the *in silico* organoid, and because the initial structure of human colon organoids is spherical (Fig. 1), we randomly deposited seeds onto a normalized sphere surface (Fig. 2B). Based on the mean number of cells in our analyzed biological organoids (Table 1), we set-up 400 seeds for our vertex organoid models. Then, a three step meshing obtained the cell volumes. A Delaunay tessellation connected neighboring seeds and a Voronoi tessellation delimited cell area into the 2.5D shell. Finally, we duplicated and connected two copies of this 2.5D shell, creating a 3D closed monolayer spherical tissue.

Once the 3D volume generated, we incremented dimensionalities, which are the radius of the spherical structure, based on the radius mean measured for each organoid stage (cystic/intermediate/columnar), and a relative tissue thickness as the ratio of the mean thickness normalized by mean radius of the biological organoids at each stage (Table 1). Thus, to mimic either the cystic/intermediate/columnar organoid stages, we incremented into our *in silico* model relative tissue thickness of respectively 0.07, 0.25 and 0.4 (Fig. 2C). These steps generated vertex organoid model for each organoid stage (Fig. 2C a, b and c).

Onto these models, we then tested active vertex models previously proposed to study global epithelial deformation^28,29^. These previous models use 2D solvers based on a simple energy functional equation (comparable to a behavior law) to model epithelial monolayer relaxation as described in Farhadifar 2D model^30^.

We generalized and applied this problem to 3D^31,32^. The mathematical formulation of this 3D Active Vertex Model (AVM)^33^ can evolve to include volume elastic coefficient for cell compressibility, face elastic coefficient and lumen compressibility (equation (1)).

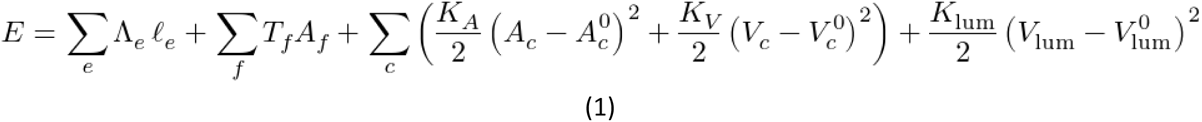

Where Λ_e_ is the line tension coefficient, le is edge length, T_f_ is the surface tension coefficient, Af is the face area, and K_A_, K_V_, K_lum_ are the elasticity coefficients for respectively the global cellular area (Ac), the cellular volume (Vc), and the luminal volume (Vlum). The index 0 corresponds to the initial reference state.

The minimum energy equilibrium is achieved by a gradient descent strategy using Broyden-Fletcher-Goldfarb-Shanno constrained minimization algorithms. We solved the system by adapting the mechanical parameters (Table 2), determined to match at best the observed biological morphologies of the organoids (Fig. 1). This led to the deformation of the vertex organoids (from Fig. 2Ca, b, c to Fig. 2Ca’, b’ and c’) *via* the vertices displacement, thus giving access to the modification of the morphology of the vertex organoids. We could then obtain information on individual cells regarding volumes, apical and basal areas, lateral areas, and aspect ratio due to the evolution of organoid morphology (Fig. 2C a’, b’ and c’).

**Table 2:** Rheology model parameters. Definition of the parameters used for the resolution of the organoid architecture equilibrium, using Tyssue quasistatic solver (AVM simulation)

| Parameter Name | Type | Value | Unit |
| --- | --- | --- | --- |
| Line tension | edge | 50.0 | pN |
| Surface tension | face | 10.0 | pN$\mu\text{m}^2$ |
| Contractility | face | 0.1 | pN $\mu\text{m}$ |
| Area elasticity | cell | 1000.0 | pN/$\mu\text{m}/\mu\text{m}^4$ |
| Volume elasticity | cell | 1000.0 | pN/$\mu\text{m}/\mu\text{m}^6$ |
| Volume elasticity | lumen | 0.001 x cell number | pN/$\mu\text{m}/\mu\text{m}^6$ |

Comparing data from the biological (Fig. 1F) and the vertex (Fig. 2C) models, we observed that this approach is not efficient and sufficient. In particular, after resolution of equation (1) in the vertex models (cystic/intermediate/columnar), we observed a uniformity in the cell volume distribution and in the cellular area. Overall, the cellular data obtained are not representative of what observed on the biological organoid model at the different stages.

### Incrementation approach of FEM onto vertex organoid model

#### Conceptual approach

This discrepancy between the two models can be explained by the fact that this modeling approach does not consider and increment the physical constraints undergone by the biological organoid model. One way to account for these constraints is to add material properties and stress information into the deformation of the vertex model *via* the finite element method (FEM). This result is achieved in two steps. Indeed, we consider that the equilibrium state of the organoid morphology depends on both the material properties related to the object to be simulated, and the constraints present in the system^34–39^. Thus, by applying first a deformation onto the vertex model into which material properties have been incremented, stress information can be extracted. Secondly, by reverse approach, FEM allows to test the relevance of this material properties/stress couple to obtain an equilibrium state of the simulated model (Fig. 3A).

**Figure 3.**
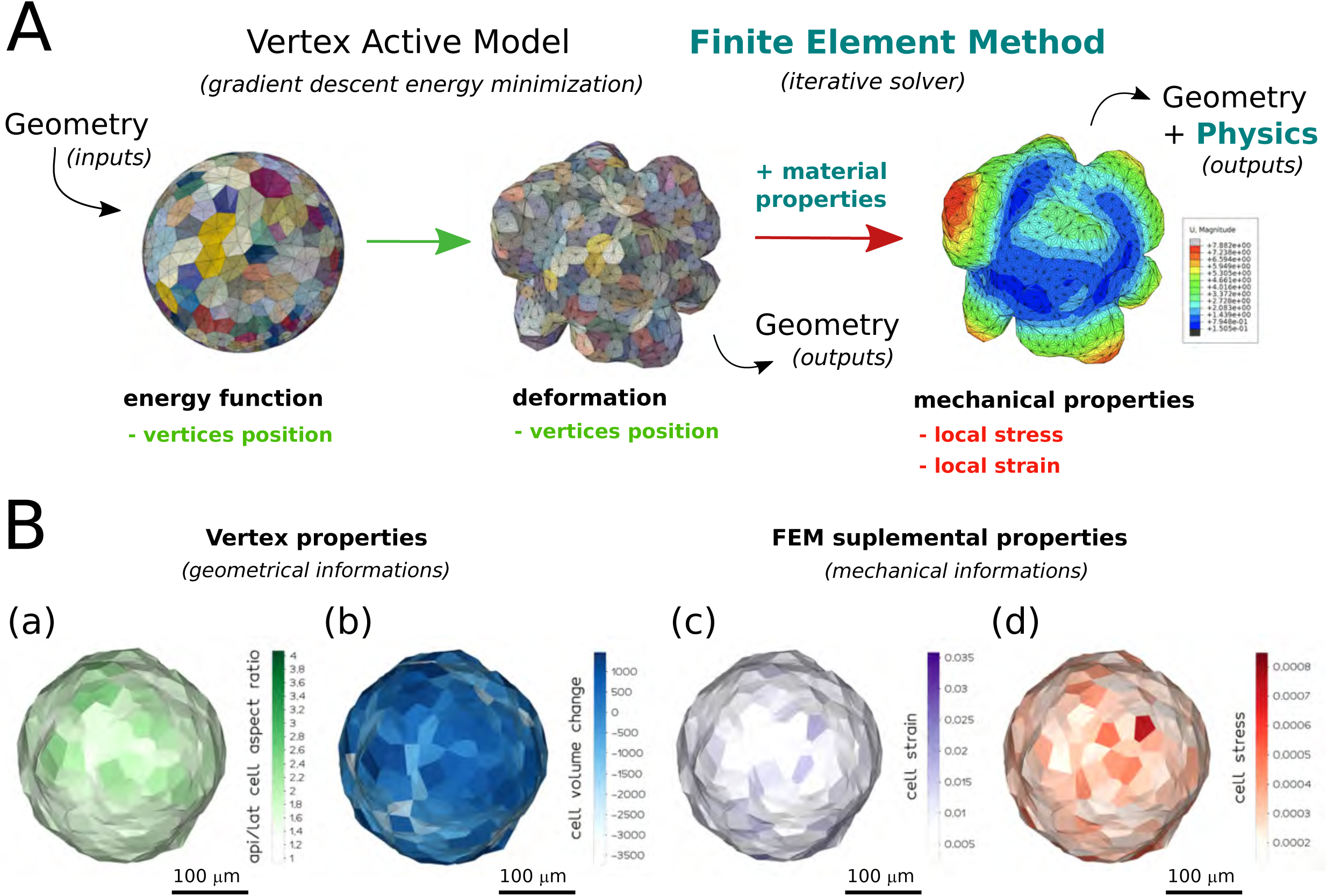
A/ Schematization of the implementation of the finite element method (FEM) on the deformation of 430 cells *in silico* organoid resolved with a 3D Active Vertex Model (AVM). In this study, we propose to add FEM analysis and material properties to study geometrics and physics relationship in organoids architecture to access mechanical properties. B/ In addition to the geometrical data like apical/lateral area cell aspect ratio (A-L cell aspect ratio, (a)) or cellular volume, here the differential volume after *versus* before deformation (cell volume change, (b)) that are mesh intrinsic properties, and accessible in all vertex models, the use of FEM allow to compute a mean strain value (c) and a mean cell stress (MISES) (d) for each cell of the organoid. Each property is measured at the element unit and averaged at the cellular unit for each 430 cells of the *in silico* organoid represented.

The interest of applying FEM onto the vertex model is that we can extract both cell stress and cell strain from the *in silico* organoid (Fig. 3B). Regarding cell stress, FEM allows to verify the von Mises^40^ yield criterion (MISES), also named ‘equivalent tensile stress’, in the post processor informing about the resulting stress for each element after deformation. Here, MISES will be used to evaluate the stress level for each individual cell, defining cell stress. Additionally, cell strain can be extracted as the mean value of the strain of each element constituting the analyzed cell, itself computed *via* FEM.

#### Modeling organoid deformation

Because our model focuses on the use of living material, we characterized the interplay between tissue and cell deformations to design our multi-scale model (from the cell morphology to the global organoid morphology). We observed three types of local epithelial morphological alteration in the columnar organoid (Fig. 4A). The first presents a tissue curvature without important cell shape or thickness changes (Fig. 4Aa). The second displays a cellular thickness decrease at local point (Fig. 4Ab), while the third exhibits an increase of the epithelial monolayer along with a local cell size increase (Fig. 4Ac).

**Figure 4.**
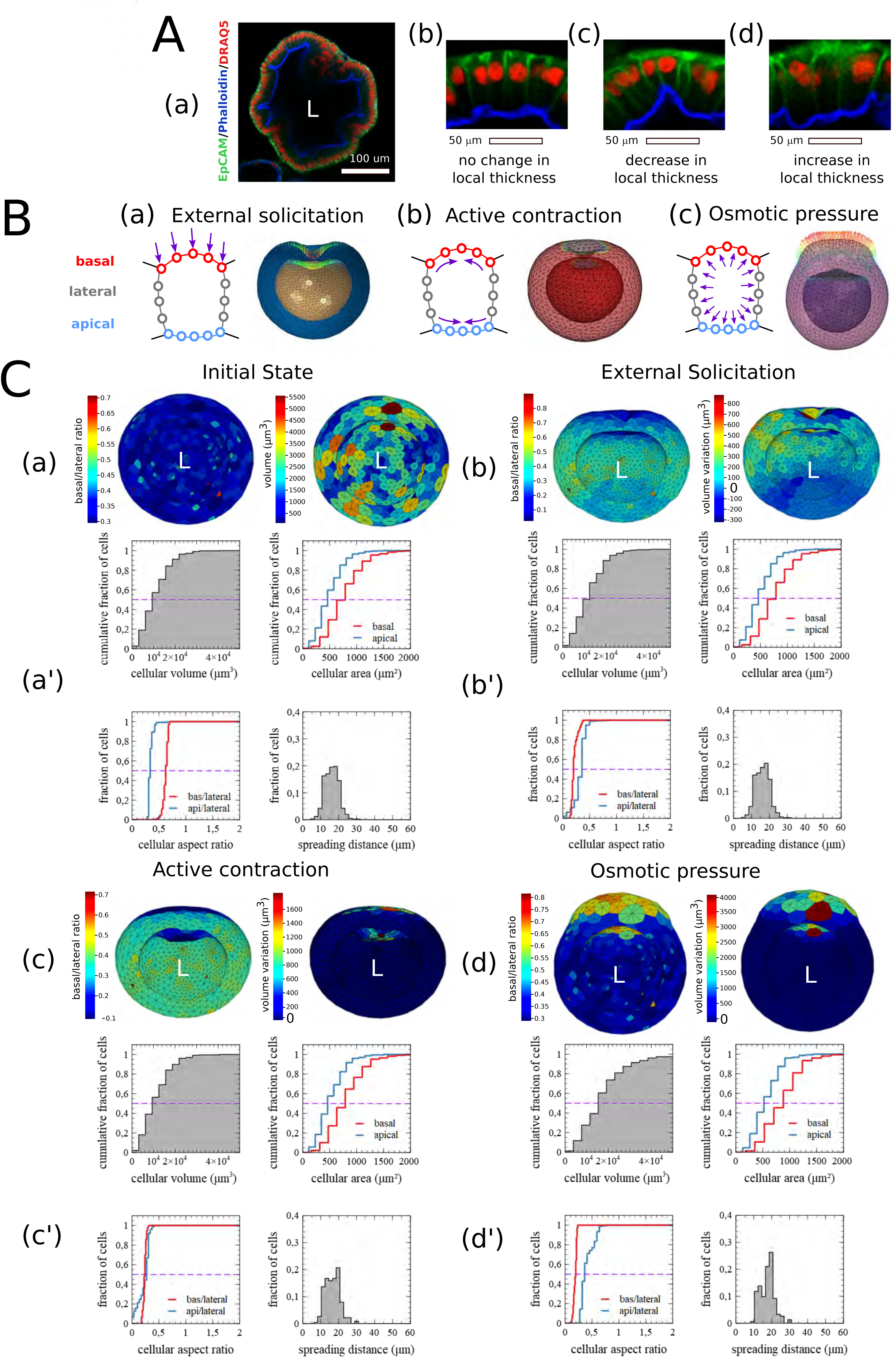
A/ (a) z section of a fixed columnar organoid with phalloidin (blue) immunostaining against actin filaments, EpCAM (green) immunostaining against epithelial cell adhesion molecules and nucleus Draq5 (red) staining. (b-d) Three zoom snapshots realized on the same fixed organoid presenting three distinct cases of deformation with no, negative or positive change in the local thickness of the epithelial monolayer. B/ Schema presenting the three different types of controlled loads tested in our FEM model. (a) external solicitation controlling node displacement, here a gradual push on the apical surface resulting in an invagination phenotype. (b) Active contraction of both apical and basal surface defining a negative relative temperature, resulting in an invagination phenotype. (c) Osmotic pressure imposed at the surface of membrane elements, positive pressure dilates the cells and drive an increase in the local thickness of the tissue. C/ Sensibility of the 3D *in silico* organoid FEM model through controlled load constraint. Het map 3D plot of two difference geometrical data, basal/lateral area cell aspect ratio and volume at the surface of the cells for a for the common initial state (a’), a case of external solicitation (b’), a case of active contraction (c’), and one case of osmotic pressure (d’). 3d representation performed using Vedo (python). Morphology description card of three different organoids resumed with four different distribution plots for the cellular volume, basal (red) and apical (blue) surface area, basal/lateral (bas/lateral, red) and apical/lateral (api/lateral, blue) aspect ratio and spreading distance distribution for the common initial state (a’), a case of external solicitation (b’), a case of active contraction (c’), and one case of osmotic pressure (d’). All the tests are using the same 3D *in silico* organoid constituted of 400 cells (a, a’).

To generate the *in silico* model, we first considered an initial perfectly spherical configuration (Fig. 2B). Using the FEA solver Abaqus©, we decomposed the shape changes by implementing three types of stresses/loads to obtain morphological elementary deformations into the *in silico* model generated based on the materials properties (Table 3 and Fig. 4B). Indeed, from a mechanical perspective, the term ‘load’ will be used to express the stress imposed to the biological model.

**Table 3:**
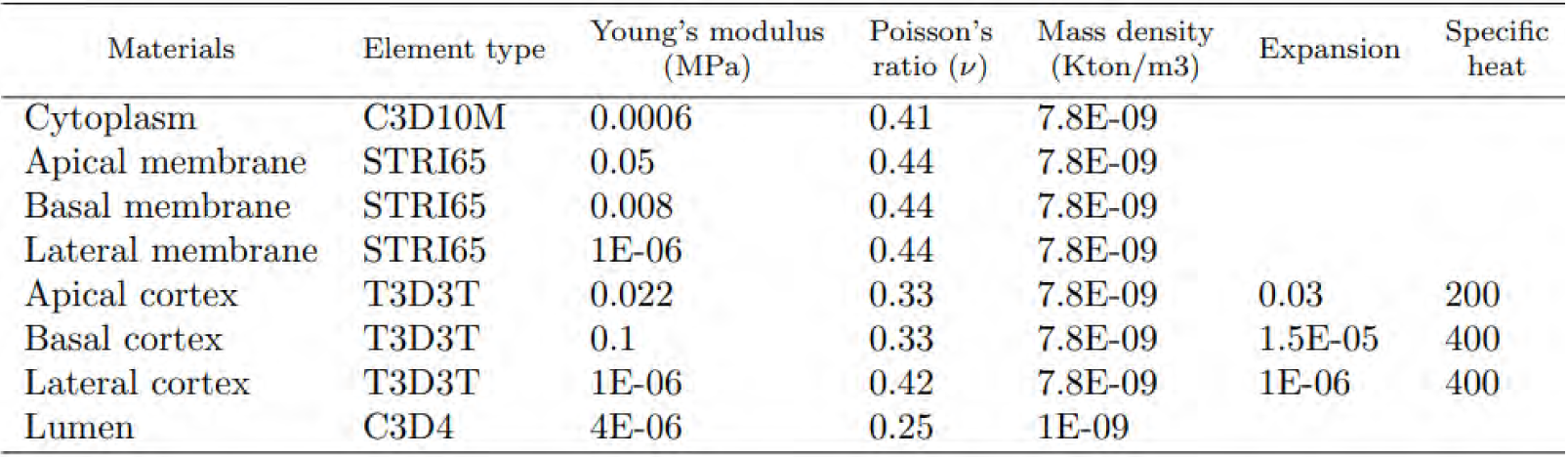
Material parameters used in the *in silico* organoid FEM model obtained by manual optimization based on both relevant global organoid morphology simulations and the literature.

| Materials | Element type | Young's modulus (MPa) | Poisson's ratio ($\nu$) | Mass density (Kton/m<sup>3</sup>) | Expansion | Specific heat |
| --- | --- | --- | --- | --- | --- | --- |
| Cytoplasm | C3D10M | 0.0006 | 0.41 | 7.8E-09 |  |  |
| Apical membrane | STRI65 | 0.05 | 0.44 | 7.8E-09 |  |  |
| Basal membrane | STRI65 | 0.008 | 0.44 | 7.8E-09 |  |  |
| Lateral membrane | STRI65 | 1E-06 | 0.44 | 7.8E-09 |  |  |
| Apical cortex | T3D3T | 0.022 | 0.33 | 7.8E-09 | 0.03 | 200 |
| Basal cortex | T3D3T | 0.1 | 0.33 | 7.8E-09 | 1.5E-05 | 400 |
| Lateral cortex | T3D3T | 1E-06 | 0.42 | 7.8E-09 | 1E-06 | 400 |
| Lumen | C3D4 | 4E-06 | 0.25 | 1E-09 |  |  |

The first load corresponds to an external solicitation resulting in surface node displacements at the cell basal side (Fig. 4Ba). Cell external forces, defined by displacement at the node level, are a common way to compute the resulting strain and constraint with FEM. We used this capability to model epithelial invagination in our FEM organoid. To mimic this local deformation, we imposed a displacement field on the basal cell surface with an inverted Poisson law distribution. The application of this positive pressure, external to the organoid, conducts to an invagination-like deformation with a curvature of the basal surface accompanied by the deformation of the apical surface (Fig. 4Cb). Because curvature measure by itself is relative to the scale of the object analyzed, we used as curvature descriptors the apical/lateral and basal/lateral cellular aspect ratios to quantify the regional organoid deformation at the cellular scale. Compared to Fig. 4Ca and a’, and as observed in Fig. 4Cb and b’, external solicitation principally affects the basal/lateral and cellular aspects ratio but has limited impact on the volume variation. Moreover, no obvious change in spreading distance was noticed.

The second load, active contraction, implies a contraction *versus* relaxation of cortical elements (Fig. 4Cc and c’). Indeed, forces at the cell top surface are linked to the need for apical contraction during the invagination of the epithelium, while forces at the bottom surface of the cell are more commonly associated with cell expansion, as the basal actin cytoskeleton is actively disassembled^41^. We choose to model the differential cortical contraction using a FEM volume expansion model which is commonly used to model thermal contraction or expansion of mechanical structures. The contraction relies on two related material parameters, the expansion rate and a specific heat. The expansion rate is the ratio between the elementary volume variation and the elementary load variation. From a mechanical perspective, the specific heat corresponds to the amount of energy needed to increase the elementary volume by one unit of load. In our perspective, it corresponds to the ability of our biological structure to inflate in response to an external load. Thus, the simulation of differential contraction at the apical and basal surfaces of the cells was done by defining two expansion rates and a temperature condition allowing deformation (Table 3). Cell expansion rates are directly linked to the activity of the actin cytoskeleton. We mimic this interplay by setting-up a higher expansion rate at the apical surface compared to the basal surface. We found that a light loading can be considered as a cellular expansion parameter to induce an invagination-like deformation of the organoid (Fig. 4Cb and b’). As a result, compared to Fig. 4Ca and a’, active contraction induces changes in apical/lateral and basal/lateral cell aspect ratios and important decrease in volumes (Fig. 4Cc and c’). Again, no obvious change in spreading distance was observed.

The third load, osmotic pressure, coincides with cytoplasmic pressures exerted onto cell surfaces, from basal to apical (Fig. 4Cd). If cytoplasmic pressure is a function of actomyosin contractility and water flow, the independent control of the last one *via* osmotic forces regulation, is another important component emerging from mechanical models^42^. In our model, we considered that cell cytoplasms are purely elastic^39^ (Table 3). We thus defined their Young’s modulus and a Poisson’s ratio which can be linked to the cellular compressibility. Using FEM, we tested changes in osmotic pressure applying pressures forces directly onto specific cell surfaces as loading parameters. Compared to Fig. 4Ca and a’, as observed in Fig. 4Cd and d’, osmotic pressures induce changes in cellular aspect ratio, an important volume increase as well as spreading increase.

As a conclusion, our FEM has been developed to model the three main loads and types of elementary deformations that can affect the tissue architecture. Indeed, applying local loads onto the tissue allows to extract cellular characteristics. Thanks to this approach, we validated the material parameters describing elasticity and volume expansion in our model. Moreover, for each of them, we identified the ranges of loads to input into the *in silico* model that are necessary to simulate morphological alteration occurring within the establishment of the organoid morphology.

### Model validation

After working at cellular level, we then tested our approach on our global *in silico* organoid. Since the biological organoids are grown in Matrigel matrix surrounding them, we could consider that the external solicitation is equivalent all around the organoid and thus either negligible in the *in silico* model. Nevertheless, it could be easily added by applying a unique scale factor on all the different loads. To simplify the modelling approach, we neglected the external load in the global *in silico* organoid model. Whether at cellular or organoid scale, the active contraction is dependent on the concentration and activity of the actin cytoskeleton. Between these two parameters, the activity is the one mainly responsible of the alteration of the morphology^43^. Thus, in order to simplify the modeling approach, we defined a homogeneous actin concentration *via* the expansion coefficient and an inhomogeneous actin activity *via* a load expressed by a temperature onto the *in silico* model. In the context of biological organoid models, it is reasonable to consider that two entities exhibit semi-permeable membranes in regard to osmotic pressure. In one hand, the cellular membrane defines an intracellular osmolarity. In another hand, the closed epithelial monolayer defines an intraluminal osmolarity. Thus, the contribution of these two osmolarities can explain the differential pressure presents at the basal/external and apical/luminal surface of the global epithelium monolayer of the organoid. This is modelized in our *in silico* model by two different surface pressures necessary to obtain a global deformation representative to the one observed in the biological model (Fig. 1C).

Based on these concepts, to understand the mechanics driving dynamical morphological changes during the live transition from a cystic to a columnar morphology in the biological organoid model, we modelized the three steps representing respectively the cystic, intermediate and columnar morphologies (Fig. 5A). In step 1, to obtain the cystic morphology, we imposed osmotic pressures of 100 Pa and 20 Pa respectively at the apical and basal surfaces of the *in silico* organoid. In step 2, to mimic the decrease of the global biological organoid size and its lumen, as well as the swelling of the apical surface (Fig. 1C), we increased the apical and lateral active contractions by lowering the volume expansion load of the actin-cytoskeleton at −100 a.u. in parallel to a relaxation of the contraction at the basal membrane by increasing the volume expansion load of the actin-cytoskeleton to 100 a.u.. In addition to the volume expansion load control, we concomitantly decreased the osmotic pressure inside the lumen with the respective osmotic pressure at the apical and basal surfaces of −30 Pa and −10 Pa. Finally (step 3), regarding the columnar organoid morphology, the increase in the cellular area and volume (Fig. 1Fc) was simulated by increasing the osmotic pressure inside the cells, consequently the swelling was adapted by increasing the active contraction of the apical actin-cytoskeleton (−200 a.u.). To support the increase of the osmotic pressure inside the cell with the respective osmotic pressure at the apical and basal surfaces of −30 Pa and −10 Pa, we then increased the relaxation of the lateral tension by in increasing the volume expansion load of the lateral actin-cytoskeleton (300 a.u.).

**Figure 5.**
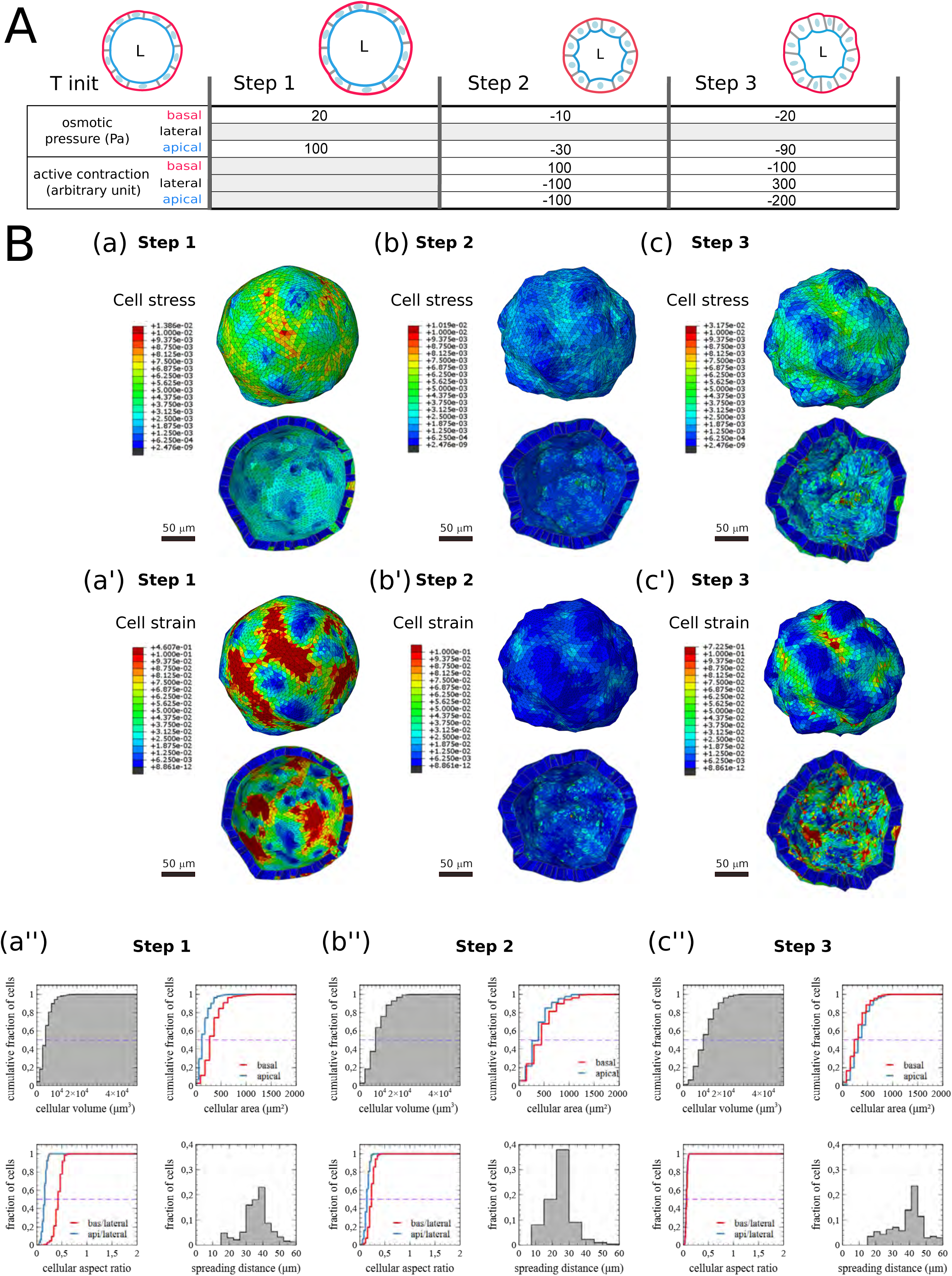
Human intestinal organoid FEM model simulation of the deformations occurring from cystic morphology to columnar morphology during polarization event. A/ Schematization of the loading sequence used in the simulation model. Consecutives osmotic pressures and active contraction are imposed to the apical (blue), basal (red) and lateral (grey) cellular surfaces of the *in silico* organoid model. B/ Results of the FEM simulation for the step1 (a, a’, a’’), the step 2 (b, b’, b’’) and the step 3 (c, c’, c’’) equivalent respectively to the cystic, intermediate, and columnar organoid morphology. Stress (MISES) plot at a subcellular resolution at the global view (basal) and cut view (apical) of the in silico organoid model (a, b, c). Strain deformation plot at a subcellular resolution at the global view (basal) and cut view (apical) of the in silico organoid model (a’, b’, c’). Morphology card of the in silico organoid model with four different plots for the cellular volume, basal (red) and apical (blue) surface area, basal/lateral (red) and apical/lateral (blue) aspect ratio and spreading distance distribution (a’’, b’’, c’’).

As a result, regarding cell stress, in step 1 (cystic morphology, Fig. 5Ba), it is distributed non-uniformly between the cells within the global organoid morphology. External view (upper panel) presenting the stress at the basal membranes displays areas with higher stress, while the internal view (lower panel), corresponding to the apical membranes, shows an overall decreased stress compared to external view. Moreover, on each view, we can observe areas with higher stress associated to higher deformation. In step 2 (intermediate morphology, Fig. 5Bb), the overall organoid is more relaxed than in step 1, certainly due to the decrease in osmotic pressure in the lumen. On the external view, no area display either cell stress or local deformation, while on the internal view, areas remain with higher stress associated to the global organoid deformation. Finally in step 3 (columnar morphology, Fig. 5Bc), the global range of cell stress is increased compared to step 2, but equivalent to the one observed in step 1. Here high cell stress intensity is only restricted to few cells, mainly accumulated at their apical side, correlating with the global organoid deformation.

Cell strain localization in step 1 (Fig. 5Ba’) correlate with the organoid global deformation. In step 2 (Fig. 5Bb’), on both views, the organoid is relaxed displaying only very low level of cell strain. Finally in step 3 (Fig. 5Bc’), while low cell strain is observed on the external view, the internal view shows that higher local cell strain is associated with global organoid deformation. Overall, in step 1 and step 2, cell strain is also non-uniformly distributed but with a strong correlation with the cell stress pattern. However, in step 3 and more specifically on the internal views, cell stress and cell strain patterns are mutually correlated. These findings are coherent as we first modulated the organoid inflation playing on the global pressures while we simulated the organoid global constriction by increasing apical contraction.

Then, as done previously, we obtained individual cell information on the volumes, apical and basal areas, lateral areas and aspect ratio due to the evolution of the organoid morphology (Fig. 5Ba’’, b’’ and c’’). Comparing results extracted here compared to the ones from the biological model, we notice that the implementation of FEM onto our vertex model allows to obtain an *in silico* organoid model displaying cellular parameters that are relevant to the biological model. Thus, thanks to this approach, we validated the material parameters describing elasticity and volume expansion in our *in silico* model. Finally, we identified the range and types of loads (external solicitation, active contraction and osmotic pressure) involved in the establishment of global organoid morphologies (cystic/intermediate/columnar).

Finally, we validated the biological relevance of our FEM-implemented vertex model using a reverse approach. We performed a forskolin-induced swelling assay. Forskolin is a CFTR activator leading to ions and water uptake into the organoids to equilibrate osmotic pressure^44^.

To induce the organoid swelling, we treated the organoids with Forskolin (Fig. 6). As expected, comparing the organoids under forskolin *versus* control condition, we observed an increase in both global organoid and individual cell volumes (Fig. 6A).We then extracted the biological data obtained (cellular volume, area, aspect ratio and spreading distance.

**Figure 6.**
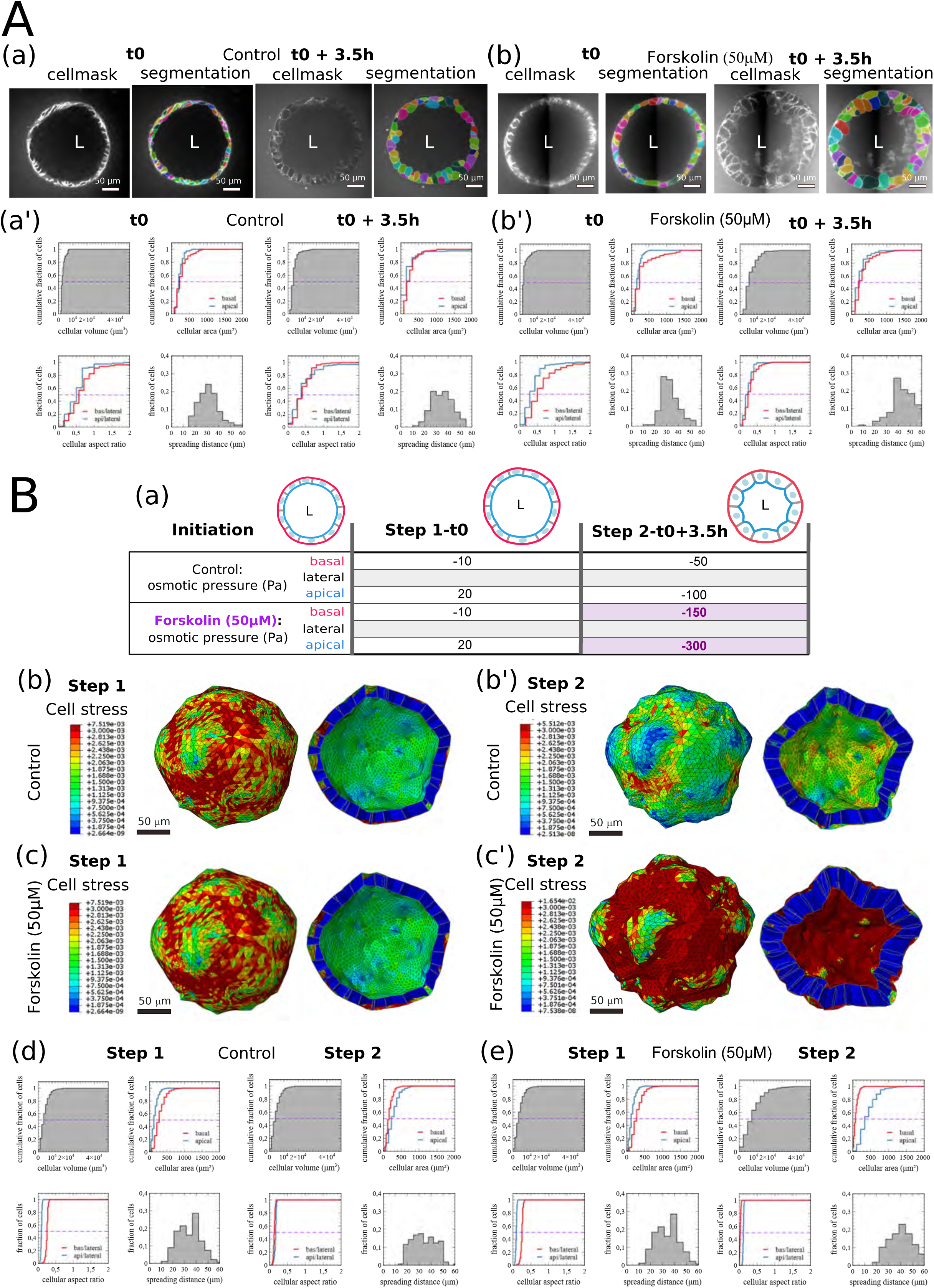
Human intestinal organoid FEM model simulation of a swelling assay using Forskolin to increase osmolarity. A/ Quantification of organoid morphology during swelling assay. Based on cellmask live label imaged on confocal microscopy, (b) cellular compartments for control organoid (a) or organoid treated with Forskolin at a 50μM concentration (b) before and 3.5 hours after treatment. Morphology description card are dressed for control (a’) or swelling condition (b’). B/ Schematization of the loading sequence used in the swelling assay model (a). Consecutives osmotic pressures are imposed to the apical (blue), basal (red) and lateral cellular surfaces of the *in silico* organoid model. Results of the FEM simulation for the step1 (b, c), the step 2 (b’, c’) equivalent respectively to the control or swelling condition. Stress (MISES) plot at a subcellular resolution on a global or cut view of the in silico organoid model (b, b’, c, c’). Morphology card of the in silico organoid model with four different plots for the cellular volume, basal (red) and apical (blue) surface area, basal/lateral (red) and apical/lateral (blue) aspect ratio and spreading distance distribution for the control (d) and the swelling assay condition (e).

To challenge our *in silico* model, we aim to mimic the observed biological organoid swelling on our FEM-implemented *in silico* organoid model. Based on the sizes and numbers of cells present in the biological organoids imaged in the swelling assay, we first generated a FEM-implemented *in silico* model of 320 cells corresponding to the modelling initiation stage. We then generated a model reproducing at best the parameters (cellular volume, area, aspect ratio and spreading distance) extracted from the biological model (Fig. 6Ba, b, c; Step 1-t0). Then, to model the effect of the swelling, we only modulated the loads corresponding to the osmotic pressure on both basal and apical surfaces (Fig. 6Ba, b, c; Step 2-T0+3.5h).

Comparing parameters from the biological models (Fig. 6Aa’ and b’) to the ones extracted from our *in silico* models (Fig. 6Bd and 6Be, respectively), we observed that the evolution of the organoid structures between the t0/step 1 and t0+3.5h/step 2 stages resulted in similar global and cellular changes. More than identifying that a three-time higher osmotic pressure load is sufficient to reproduce the forskolin-induced swelling, FEM allowed to access the evolution of cell stress among the organoid structures (Fig. 6Bb and c).

The observed correspondences between the geometrical descriptors (cellular volume, area, aspect ratio and spreading distance) of both the biological and *in silico* models represent an experimental validation of the biological relevance of our FEM-implemented vertex organoid model.

Thus, we identified and incorporated mechanics *via* the use of materials properties into our *in silico* organoid FEM model. This model recapitulates the evolution of global organoid morphology in a biologically relevant way while considering the individual cellular specificities. Taking into consideration the three main stresses/loads that can affect tissue architecture and, applying these stresses locally into the *in silico* tissue, allowed us to decipher the mechanical cellular properties involved in the establishment of the organoid morphology.

## DISCUSSION

The human colon organoid we chose as biological organoid model to study the interplay between biology and mechanics involved in tissue architecture establishment allowed to extract, by imaging approach and analysis, cellular volumetric measurements. We then calculated morphological properties (cellular area, cellular volume, cellular aspects ratio and spreading distance) characterizing the biological organoid model. These data, obtained for the cystic/intermediate/columnar morphological stages, were evaluated as possible quantitative geometrical descriptors of tissue architecture evolution during the organoid morphogenesis. Such quantitative descriptors could eventually serve as discriminant parameters to implement into clustering analysis based on tissue morphology survey by machine learning approach.

Previously, three major categories of *in silico* epithelium models depending on their structure and usage, have been proposed to study epithelial tissue behavior. Structured grid models, ideal to study self-organization of the tissue with the diffusion of chemical gradients^45^, modelized cells by the repetition of equivalent and uniform elements that makes difficult to consider cell shape variability. By contrast, unstructured grid models, which do not use predefined cell shape, are more suitable for morphogenesis studies and have been extensively used for mechanical studies on 2D tissue organization^13,30,46,47^. Agent Based Models (ABM) uses discrete tissue organization, with autonomous agents (spheres) representing the cells, to decompose the general problem, such as cell migration, cell differentiation or growth rate, in a sum of individual problem^34,48^. ABM are powerful for tissue patterning but the discontinuity in the geometrical structure makes it difficult to use for mechanical properties studies^49^.

Here, to consider both biological and mechanical properties involved in the organoid morphology evolution in a 3D multi-scale approach, from the cell to the tissue, we adapted an unstructured grid model, the 3D Active Vertex Model (AVM) describing an epithelial monolayer theoretical biophysical 3D model^20^. However, comparing the quantitative descriptors values given by this adaptation to the one extracted from the biological model, the AVM is not sufficient (Fig. 2). This discrepancy between the biological and vertex models can be explained by the fact that this approach does not consider and increment the physical constraints undergone by the biological model. One way to account for these constraints is to add material properties and stress information into the deformation of the vertex organoid model *via* the finite element method (FEM). The FEM main advantages are to allow access to a finer level of simulation detail, to integrate complex behavioral laws and to preserve the volume coherence of the cells. The similarities observed between the data of the biological model and those of the FEM model allow us to validate our model geometry (number of nodes, links, interface…), its mechanical properties (active contraction and osmotic pressure) and the imposed loads (external solicitation). Thus, the consistency of our simulation with the observed morphological alterations, and their distribution and amplitudes validate the choice of our FEM model and its use.

FEM is well suited to increment specific local mechanical characteristics into the model. Indeed, vertex model only defines vertex, faces and vertices to access the tissue. FEM allows to increment and simulate more details into any continuous models, and is thus very suitable to model a tissue. Another advantage is its multi-scale capability, allowing an interplay between the FEM and cellular elements (Fig. 2A), aka quadratic tetrahedral element & cytoplasm, quadratic shell element & membrane, quadratic truss element & cytoskeleton), but also with the tissue behavior (external solicitation, active contraction and osmotic pressure). Moreover, FEM based on material properties definition permits to fit at best the biological reality, since FEM allows to keep a volumic coherence of each individual cell. Furthermore, we introduce a volumic expansion factor to the individual cell within the model to pilot each individual cellular deformation and thus allows to model the global deformation of the tissue in biologically relevant range (Fig. 5 and 6).

Our innovative FEM model takes advantage of the accurate description of each cell and specific mechanical laws related to cell behavior to model the global tissue. This multi-scale model is a promising tool for the comprehension of global tissue alteration, and a possible contribution of individual cellular events. A reverse approach, from deformation observation to stress and strain evaluation, could help evaluating material properties evolution at cellular scale. From a biological and medical perspectives, this capacity to link individual cell behavior to global tissue alteration could find future applications in image-based diagnostic and screening approaches. Indeed, incrementation of our *in silico* FEM model into novel AI driven automatic processes could help identifying very early tissue alteration in patient’s colon organoid and provide a better personalized follow-up.

## MATERIAL AND METHODS

### Human samples

Colon samples were obtained from biopsies of patients undergoing endoscopy at the Toulouse University Hospital. Patients gave informed consent and were included in the registered BioDIGE protocol, approved by the national ethics committee (NCT02874365) and financially supported by the Toulouse University Hospital.

### Organoid culture

Colorectal crypt isolation was performed as described previously54. Fresh Matrigel (Corning, 356255) was added to isolated crypts. 25µL of Matrigel containing 50 crypts were plated in each well of a pre-warmed 8-well chamber (Ibidi, 80841). After 3 days of cultures, organoids were frozen to constitute a stock. From this stock, organoid were thawed, plated in Matrigel and expanded for amplification in culture conditions described previously^50^. After 10 days of cultures, medium was replaced by 250µL of Intesticult ODM human basal medium (StemCell technologies, 100-0214), allowing generation of intestinal organoid cultures with physiological representations of the stem cell and differentiated population. Cultures were followed for the time indicated in Figures. In order to perform the swelling assay, cultures were treated with Forskolin (50µM) or control conditions (DMSO, 1.25%) and imaged just before adding the drug (t0) and 3.5h post treatment (t0+3.5h).

### Immunostaining and live imaging

Organoids were fixed with a PBS solution containing 3.7% of formaldehyde solution for 5 min at 37°C. Then, organoids were washed in PBS and permeabilized with a PBS solution containing 0.5% Triton X-100 for 20 min at 37°C. After washes, DNA was stained with Draq5 (Ozyme, 424101, 2 µM) for 30 min, membranes were stained using anti-EpCAM antibody (Cell Signaling Technology, VU-1D9) and actin network using Alexa Fluor 594 phalloidin (ThermoFisher, A12381, 40nM). Finally, cover glasses with Matrigel domes were immersed in PBS for imaging. For live experiments, cytoskeleton was stained with SPY650-FastAct or BioTracker-488 microtubule dye (SCT142, Sigma-Aldrich), DNA was stained with the SPY555-DNA probe (SpyDNA). All these fluorescent live cell probes come from Spirochrome and were used at a 1/500 final dilution 4 hour before live start. Image acquisitions were performed using a two-photon microscope (Bruker 2P+, 20X diving lens objective) for fixed experiments and with the Opera Phenix HCS microscope (Perkin Elmer, 40X objective) for live experiments. Images were analyzed using the ImageJ software from FiJi^51^.

### Image analysis

Cell segmentation was performed using Cellpose2 machine learning based 2D approach adapted for each individual organoid analyzed and then 2D segmentations were stitched in 3D using the same software^25^. Luminal volume is segmented manually using Napari^52^ software. 3D isometric interpolation was performed using VT python libraries^53^ for segmented images (labels) or using Napari for rawdata and manual corrections of the segmentation was performed using Morphonet^54^ or Napari to select and correct only cells present entirely in the 3D acquired field in order to have correct estimation of the cellular interfaces. The number of the cells selected for the analysis is presented in Table1.

### Model Implementation and data analysis

We used Python (3.7.11) libraries NUMPY and SCIPY to generate the mesh staked in Pandas datasets that can be exported in HDF5, OBJ (for surfaces) or VTK formats using MESHIO. A large part of the python codes is using TYSSUE^33^ library for monolayer creation. For remeshing and properties extraction, CGAL and VT^53^ libraries containing image oriented algorithms written in C are used as well as VTK or Vedo^55^ for the visualization. A conversion into Abaqus© (Dassault Systems) input file was code in python to create our definitive *in silico* organoid.

#### Finite element formulation

A collection of elements convenient for our model and preconfigured in Abaqus© was selected. Each type of element is specific of the subcellular unit they have to mesh.

- Cytoskeleton: quadratic C3D10M Abaqus© tetrahedral element connecting the centroid of the cell and a part of the face of the cell. Tetrahedral elements was chosen, more convenient than hexahedral elements to cover uniformly a sphere^50^ and more flexible for complex cell shapes^56^.
- Membranes: quadratic STRI65 Abaqus© shell elements connecting only the nodes of the preceding volumic elements present at the surface of the cells.
- Actin cytoskeleton: the active network present in the intracellular space close to the membrane are modelized sharing the node of the membrane with first order T3D3T Abaqus© thermal truss elements.

### Model material parameters

We defined the physical behavior of the FEM organoid in four distinct domains, cell cytoplasm, cell membrane, cell cortex, and the extracellular lumen, each using specific elements, e.i. specific mechanical properties (Table3). For simplicity, it is assumed that cell cytoplasms are purely elastic^39^ and this elasticity is isotropic. In the literature, epithelial cell cytoplasm elasticity is characterized with a Young’s modulus ranging from 0.3 to 100 kPa depending on the biological model and the measurement method^36,38,39,48^. For cytoplasm elasticity in all our models, we fixed an arbitrary value of 2 kPa, after validation of simple sheets deformation, associated with a Poisson’s ratio of 0.44. Indeed, high Poisson’s ratio (near 0.5) are largely used for deformations which occurs at constant volume. Similarly, we gave an elastic behavior to the membrane elements in order to represent membrane elasticity based on the literature. Due to the different nature of the interface, we give a higher elasticity to the apical membrane compared to the basal membrane which is surrounding by a basal lamina.

The last surface of the cell, lateral membrane, at the interface between neighboring cells, has the particularity to connect two cellular membranes. In our continuous vertex model, we do not duplicate membranes for cell-cell contacts in order to simplify the problem. With this unique interface, we reduced the number of contacts to attribute and compute during solving iterative operations. The constraints at this lateral membrane interface are neglected using a low Young’s modulus of 4.5E-06 kPa. By this way, the constraint at cell-cell interfaces is mostly due to pressure equilibrium exercise by cytoplasmic pressures.

### Model loading parameters

To control deformation in our in silico organoid FEM model, we used three different types of loads.

*External solicitation* is modelized using node displacement on the basal membrane. This can be also used to studied displacement extracted from tracking information given by live analysis.

*Active contraction* is modelized using volume expansion method in Abaqus. This required of temperature displacement specific solver that is based on specific material properties, thermal expansion coefficient and specific heat, in addition to the definition of both initial and final temperature as a boundary condition. The load for active contraction is used as relative temperature decorrelated from the real temperature fixed in our biological system.

*Osmotic pressure* is modelized using a uniformly distributed pressure at the surface of the cells. We oriented this pressure using vectors directed outside the cells and perpendicular to the element area.

## AUTHOR CONTRIBUTIONS

AF and FBu designed the project. JL, AF, FBu and SS contributed to the conception of the original idea. JL developed the theory and the finite element model, performed simulations, analyzed data sets and results. EM coordinates the BioDIGE protocol, screened the patients, performed the endoscopy and biopsies; BioDIGE is supported by Toulouse University Hospital. DH and MQ established the organoid cultures; DM and LM performed organoid cultures, immunofluorescence staining, images acquisition and images analyses; AF developed the theory and its connection to the organoid model; AF, SS, VV and FBu developed the theory and did the interpretation of the results. JL, SS, FBu and AF wrote the manuscript. All authors contributed to the discussion and interpretation of the main results.

## ACKNOWLEGEMENTS

This collaborative work was funded by two Cancer Plan projects: Mocassin - Biosystem 2017, granted to AF and Melchior - MIC 2020, granted to FBu. JL is supported by both. DM is supported by Plan cancer Melchior-MIC 2020. LM is supported by the INSERM and the Région Occitanie. DH is supported by Université Paul Sabatier and the Région Occitanie. We also wish to thank the patients who agree in giving their tissue for research purpose and allowed this study, Guillaume Gay who allows us to use his Tyssue python library as well as Gaëlle Recher and Camille Douillet for critical reading of the manuscript and their suggestions.

## COMPETING INTERESTS

The authors have no relevant financial or non-financial interests to disclose. The authors have no conflicts of interest to declare that are relevant to the content of this article.

## Notes

### Competing Interest Statement

The authors have declared no competing interest.

